## Supplementary figures and images for "Zebrafish genetic model of neuromuscular degeneration associated with Atrogin-1 expression"

### MovieS

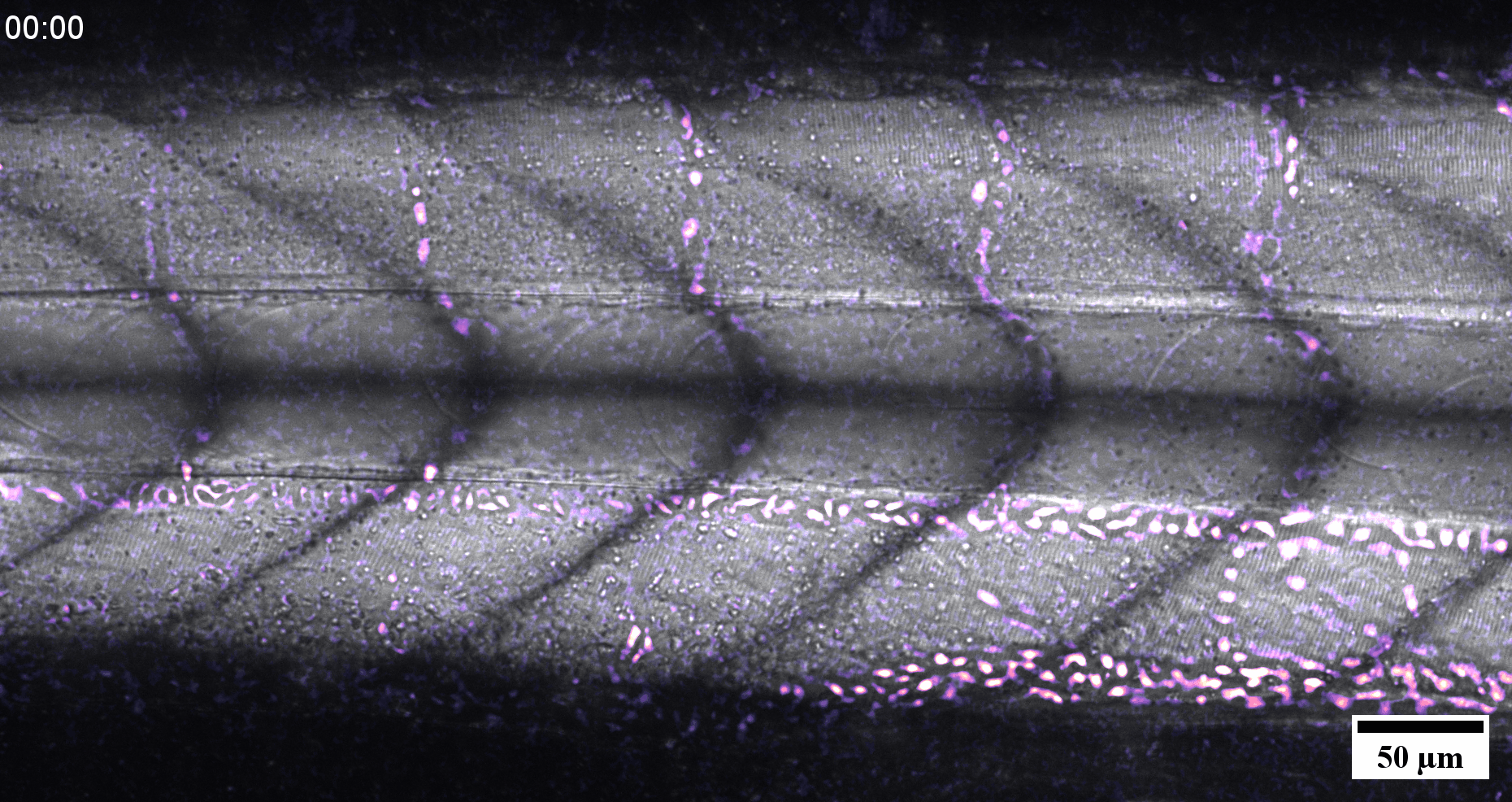
